# High sampling effectiveness for non-bee pollinators using vane traps in both open and wooded habitats

**DOI:** 10.1101/556498

**Authors:** Mark A. Hall, Eliette L. Reboud

## Abstract

1. Non-bee insects are important for pollination, yet few studies have assessed the effectiveness of sampling these taxa using low cost passive techniques, such as coloured vane traps, among different habitat types.
2. This study sampled 192 sites—108 in wooded and 84 in open habitats within an agricultural region of southern Australia. Pairs of blue and yellow vane traps were placed at each site for a period of seven days during the austral spring.
3. Overall, 3114 flies (Diptera) from 19 families and 528 wasps (non-bee and non-formicid Hymenoptera) from 16 families were collected during the study. This sampling was representative of the region, with vane traps equally or more likely to collect as many families from both taxa as those reported on the Atlas of Living Australia (ALA) database for the sampling area.
4. Blue vane taps (BVTs) had greater average richness of both flies and wasps and greater abundance of individuals than yellow vane traps (YVTs). BVTs were particularly favoured by certain fly and wasp families known to pollinate flowers (e.g. Syrphidae, Bombyliidae and Scoliidae), whilst YVTs sampled some less common fly families, such as Acroceridae and Bibionidae that also provide additional ecosystem services to pollination.
5. Vane traps are an effective passive sampling technique for non-bee pollinators, such as flies and wasps. This study supports the use of vane traps as a component of the sampling protocol for ecological census and population monitoring within multiple habitat types, to effectively sample a more complete pollinator community.

## INTRODUCTION

Understanding the distribution of bees within agricultural landscapes has received much of the research interest in pollination and community ecology (Kremen *et al*., 2002; Kennedy *et al*., 2013; Koh *et al*., 2016). This is largely due to their important role in the pollination of many native plant and crop species (Heard, 1999; Morse & Calderone, 2000; Klein *et al*., 2006). However, there is also growing evidence of the importance of non-bee insect taxa in pollination, such as flies, wasps, beetles and butterflies (Potts *et al*., 2016; Rader *et al*., 2016; Ollerton, 2017). Some of these non-bee taxa also play a vital role in other ecosystem services, such as pest control and nutrient cycling (Zhang *et al*., 2007; McCravy, 2018), and may forage more effectively than bees under different climatic conditions (Inouye & Pyke, 1988; Lefebvre *et al*., 2018). Non-bee pollinators may even replace bees as the dominant flower visitors in heavily modified environments (Stavert *et al*., 2018). Further, pollination may be more effectively achieved if multiple species, including non-bee pollinators, visit flowers (Brittain *et al*., 2013; Alomar *et al*., 2018; Winfree *et al*., 2018; Thomson, 2019). Yet to date, non-bee pollinators have been under-represented in pollination studies and there is no consensus on a passive survey method that effectively samples both the bee and non-bee pollinator fauna of a given area.

Active sampling is often used for targeting particular species, recording plant-pollinator interactions or obtaining density estimates for a given area (Larsson & Franzen, 2008; Campbell *et al*., 2016; Taki *et al*. 2018). However, sampling in this way can be time consuming, and results may be biased by the skills and experience of surveyors (Westphal *et al*., 2008; McCravy, 2018). Alternatively, passive sampling allows for a greater range of data collection (i.e. day and night, across seasons) at a relatively low cost (Saunders & Luck, 2013; McCravy, 2018). Passive sampling methods are usually easy to install and do not require any specialist skills from the operator (Missa *et al*., 2009; Westphal *et al*., 2008; Saunders & Luck, 2013). Many different trapping methods are potentially suited to sampling insect fauna, depending on the type of question being asked. A simple and effective sampling technique is the use of coloured components, which are known to attract many pollinator groups (Kirk, 1984; Pickering & Stock, 2003; Desouhant *et al*., 2010; Vrdoljak & Samways, 2012). For example, coloured pan traps are a widely used and often highly successful method for sampling the pollinator community (Westphal *et al*., 2008; Saunders & Luck, 2013). Coloured sticky traps have also been used for arthropod studies, however these can vary greatly in their ability to collect certain groups (e.g. Hymenoptera, Coleoptera, Hoback *et al*., 1999; Pickering & Stock, 2003) and make identification of small specimens difficult (i.e. <4 mm, Pickering & Stock, 2003). Coloured Malaise traps are rarely used for pollinator-specific surveys and are not as efficient as pan traps (Campbell & Hanula, 2007). However, non-coloured Malaise traps are widely used and effective for general arthropod sampling (Kitching *et al*., 2001; Campbell & Hanula, 2007; Missa *et al*., 2009). Coloured vane traps are much less studied, but have recently shown great potential, especially for bees (Gibbs *et al*., 2017; Hall, 2018). Yet very little is known about their effectiveness in sampling the non-bee pollinator community.

The quality of colours used for passive trapping varies according to trap type. When sampling pollinators using pan, sticky or vane traps, the most commonly used colours are white, yellow and blue (Abrahamczyk *et al*., 2010; Vrdoljak & Samways, 2012; McCravy, 2018). Blue and yellow coloured pan traps are often found to be the most attractive to insects, but there is no consensus on the greater attraction of either colour for different taxa or in varied habitat types (Kirk, 1984; Campbell & Hanula, 2007; Abrahamczyk *et al*., 2010; Saunders & Luck, 2013). This is possibly due to variance in reflectance and vibrancy owing to a lack of consistent quality of colour and design (e.g. pre-painted bowls: Campbell & Hanula, 2007; Joshi *et al*., 2015, self-painted bowls: Westphal *et al*., 2008; Shrestha *et al*., 2019). This makes comparing different pan trap studies problematic. Some pollinating insect taxa, such as butterflies and moths (Lepidoptera), also appear to be largely underrepresented in pan trap studies (Missa *et al*., 2009; Vrdoljak & Samways, 2012) but are sometimes caught in large numbers using Malaise traps (Campbell & Hanula, 2007). However, Malaise traps do not appear to be particularly efficient at sampling non-*Apis* bees (Bartholomew & Prowell, 2005; Campbell & Hanula, 2007; Missa *et al*., 2009), thus do not provide comprehensive sampling for pollinators. Alternatively, vane traps come in only two colours (yellow and blue) and their commercial availability (and thus consistency of colour) potentially allows for greater comparison between studies (Hall, 2018).

Vane traps are very effective at attracting bees without the need for pheromones or other liquids (Stephen & Rao, 2005; Hall, 2018), and are increasingly being used to sample wild bee populations worldwide (Kimoto *et al*., 2012; Lentini *et al*., 2012; Gibbs *et al*., 2017). Both Stephen & Rao (2005) and Hall (2018) found blue vane traps (BVTs) to be particularly effective at sampling wild bee diversity compared to yellow vane traps (YVTs). Yet, in the Stephen & Rao (2005) study, very few non-bee pollinators were sampled, and no reports of non-bee taxa appear in other vane trap studies, other than to mention that they were removed (e.g. Joshi *et al*., 2015).

However, because bees are often poorly represented using other passive methods typically used to survey arthropods (e.g. sticky traps: Pickering & Stock, 2003, Malaise traps: Missa *et al*., 2009), if we are to effectively sample both bee and non-bee communities, a more representative passive survey method is required. Vane traps are one such possibility as they are easy to install, provide consistent sampling effectiveness over large areas and are useable over longer time periods, compared with other widely used methods (Stephen & Rao, 2007; Kimoto *et al*., 2012; Lentini *et al*., 2012). However, to truly test the effectiveness of this trapping technique, we also need to compare results to what is known for a given area. This can be achieved by utilising online biological repositories (Belbin & Williams, 2016).

This study tests the efficacy of BVTs and YVTs in sampling the non-bee fauna of both open and wooded areas within an agricultural landscape of southern Australia. To test this, the following questions were asked:

1. Do vane traps capture a representative sample of the non-bee pollinating fauna of agricultural landscapes?
2. Is there a difference in the effectiveness of vane traps when sampling flies and wasps using different colours and/or within open and wooded environments?
3. Is there a consistent response between the two taxonomic groups and/or families within them?

## MATERIALS AND METHODS

### Study area and system

This study was conducted across a rural landscape of 11,550 km^2^ on the riverine plains of north-central Victoria, Australia, with a sampling area of ~3,600 km^2^ (Fig. S1a). The region has been transformed by human land-use, predominantly agricultural production, experiencing extensive clearing of over 80% of native vegetation (Environmental Conservation Council, 2001). Much of the wooded vegetation now occurs along linear roadsides and streams (Hall *et al*., 2018), or as isolated trees within farm paddocks (Gibbons *et al*., 2008). Human land-use consists of sheep and cattle grazing along with the cropping of wheat, oats, canola and legumes, often on a rotational basis (Bell & Moore, 2012). Average annual rainfall for the district is 492-648 mm per year, and daytime maximum spring temperatures are typically 16-27°C (Bureau of Meteorology, 2016).

Surveys were conducted within 24 individual farming landscapes (each 1 km diameter) across the study region, each containing both wooded cover and open farmland with a range in tree cover from ~5-22%, which is indicative of the region.

All sites within farms (n=8 per farm, n=192 in total) were selected such that they were similar in vegetation characteristics and topography. Major roads and river systems were avoided for consistency. For full details of methods, see Hall (2018).

### Sampling

#### Field sampling

Non-bee pollinator insects (flies and wasps) were sampled using vane traps (SpringStar^TM^ LLC, www.springstar.net), comprising a 64 oz (1892 ml) plastic collecting jar, screw top funnel and two interconnecting ultraviolet semitransparent polypropylene vanes, coloured either blue or yellow (Figs. S1b and S1c). These are known to be highly reflective when exposed to 365 nm filtered (UV-A), and midrange 302 nm filtered light (UV-B) (Stephen & Rao, 2005). Two pairs of traps (one blue and one yellow) were hung from a tree branch or pole at a height of approximately 2 m within the typical features present in the landscape. Thus, 2 pairs were deployed in roadsides with or without tree cover, 2 in creeks with or without tree cover, 2 in scattered trees within farmland and 2 in open farmland: n=108 wooded sites, n=84 open sites (see Hall *et al*., 2019 for description of landscape-scale design, and Hall, 2018 for full methods). Traps remained in place for seven days during the southern hemisphere spring months, from October to November 2015. No pheromones, liquids or killing agents were used in traps. Samples were collected after one week and stored in 70% ethanol then pinned for identification. Identification of flies and wasps to family-level was conducted using available keys (Marshall, 2017; CSIRO, 2018). Other insect taxa sampled were removed due to low sampling (e.g. Lepidoptera) or because they (e.g. Coleoptera, Formicidae) were more likely caught in traps during mass flight events related to breeding (Sullivan, 1981; McHugh & Liebherr, 2009).

#### Database search

To determine if vane traps were effective at sampling a representative fly and wasp community within agriculturally dominated landscapes containing both wooded and open habitats, we compared our sampled fauna to families known to occur within the study region. To do this, we generated an area report using the *tools* dropdown menu in the Atlas of Living Australia (ALA) spatial portal (Atlas of living Australia, 2019; https://spatial.ala.org.au/), to incorporate a comparable area that encapsulated our sampling area (~3,600 km^2^). We used only spatially valid records (i.e. those where geographic location was provided). We then compared families recorded on the ALA to those sampled within our study. Note that data provided in an area report could conceivably cover a timespan from 1600 - 2018 AD, however no survey date was provided for records with the report, so we must assume that most records span a considerable time period (likely 1900 – present), much longer than our two months of sampling conducted in a single season. To investigate if non-bee pollinators in our sampling area were also indicative of the whole region, we conducted a wider search on the ALA (11,500 km^2^) which also comprised forested and mountainous areas that were not sampled in our study (Table S2, Fig. S2). Further, species utilise resources at different times of the year, leading to substantial temporal turnover in insect communities within an area (Lambkin *et al*., 2011; Thomsen *et al*., 2016; Winfree *et al*., 2018). Thus, to help interpret differences in peak activity periods for each fly and wasp family (and thus likelihood of capture during our sampling period), records were taken from the ALA database for the state of Victoria, grouped by month in which individuals were reported on the database (Fig. S3).

### Statistical analyses

We conducted three separate analyses corresponding to our three main questions. First, to determine if vane traps were effective at sampling the non-bee fauna compared with historical records for these groups, we ran generalised linear models in package *nlme* (Pinheiro *et al*., 2018), assuming a binomial distribution for presence/absence data. We compared the number of fly and wasp families separately to those recorded on the ALA database. We ran separate models comparing presence of families (pooled richness) in different coloured traps (blue and yellow), and within different habitat types (open and wooded). Sites containing tree cover (roads and creeks with tree cover and scattered tree sites) were classified as ‘wooded’, whilst sites that lacked tree cover (roads and creeks without tree cover and open farmland) were classified as ‘open’. Overdispersion was checked using Pearson residuals (Zuur *et al*., 2013) and none was detected in any model. For one model (wasp-colour preference), we fitted a Firth’s bias-reduced logistic regression in package *logistf* (Heinze *et al*., 2018) due to BVTs containing all wasp families sampled (separation problem, Heinze & Schemper, 2002). To determine the level of overlap in families between the ALA, BVTs and YVTs, we calculated the Morista-Horn index in package *divo* (Pietrzak *et al*., 2016). This quantifies the level of overlap in families between sampling methods, giving a value range between zero (no overlap) and one (perfect overlap).

Second, to determine if there was a difference in the effectiveness of vane traps when sampling flies and wasps using different colours and/or within open and wooded environments, we ran generalised linear mixed models in package *nlme* (Pinheiro *et al*., 2018). We ran separate models to test for differences in the richness and abundance of ‘all individuals’, ‘all fly families’ and ‘all wasp families’. We assumed a Poisson distribution for richness measures and a negative binomial distribution for abundance data. Due to clustering of sites within 1 km diameter landscapes and pairing of different colour traps at each survey point, *landscape* and the survey location (*pair*) were used as random effects in the models.

Last, to test if the response of different fly and wasp families were consistent, we tested differences in the abundance of each family with >30 individuals sampled (see Table S2) between trap colours and habitat types, using generalised linear mixed models in the *nlme* package (Pinheiro *et al*., 2018). We assumed a Poisson distribution for all models, except for where overdispersion was detected (Syrphidae, Tachinidae), where we assigned a negative binomial distribution (Zuur *et al*., 2013), using the *MASS* package (Venables & Ripley, 2002). Again, due to clustering of sites, *landscape* and the survey location (*pair*) were used as random effects in the models. All statistical analyses were conducted in R (v.3.5.1, R Core Team, 2018).

## RESULTS

Overall, 3114 flies (Diptera) and 528 wasps (non-bee and non-formicid Hymenoptera) were collected during the study, comprising 19 families of flies and 16 families of wasps (Table S1). Syrphidae was the most abundant family sampled with a total of 1396 individuals caught (45 % of the total abundance), including 87 % in BVTs.

For both orders, BVTs caught more insects with 67.5 % of the total abundance (68 % for flies and 63 % for wasps, Table S1). Four fly families were found only in YVTs (Acroceridae, Bibionidae, Conopidae, Scatopsidae) whereas six fly or wasp families were captured only in BVTs (Diptera: Chironomidae, Sarcophagidae; Hymenoptera: Crabronidae, Gasteruptiidae, Mutillidae, Vespidae). Almost all of these families (only found in one colour) were sampled by a single individual (except Scatopsidae n=2, Mutillidae n=2, Vespidae n=6; Table S1).

In total, 1935 insects were caught in open areas, which represented 53% of the total abundance. Six families were only sampled in open habitat (Table S1). These open-associated families often only had one or two individuals (singletons or doubletons), however two families were found in greater numbers—Sphecidae (n=29) and Vespidae (n=6). Similarly, 14 families were only found in wooded habitats, with only five families sampled having more than one or two individuals— Lauxaniidae (n=3), Mycetophilidae (n=7), Chrysididae (n=7), Pompilidae (n=5) and Scelionidae (n=4) (Table S1).

### Effectiveness of sampling using vane traps

A total of 19 fly and 9 wasp families were recorded on the ALA within our sampling area (Fig. 1). Combined BVTs and YVTs sampled 19 families of flies, 12 of which were not reported on the ALA (one was caught only in BVTs, four only in YVTs and seven in both BVTs and YVTs). Twelve fly families reported on the ALA were not caught in vane traps during the study (Fig. 1). Overall, there was no difference in the number of fly families sampled in our study in both BVTs (Est = −0.52, SE = 0.51, P=0.31) and YVTs (Est = −0.26, SE = 0.52, P=0.61), or within wooded habitats (Est = −0.26, SE = 0.52, P=0.61; Figs. 1a, 1b). In contrast, fewer fly families were collected in open habitats compared with historical ALA records for the sample area (Est = −1.06, SE = 0.53, P=0.04; Fig.1b). There was some overlap in families recorded on the ALA and those sampled using BVTs and YVTs (flies: 30-40%, wasps: ~55%, Figs. 1a, 1c). There was greater overlap based on habitat for wasps (~66%) than flies (20-35%) (Figs. 1b, 1d).

**Figure 1:**
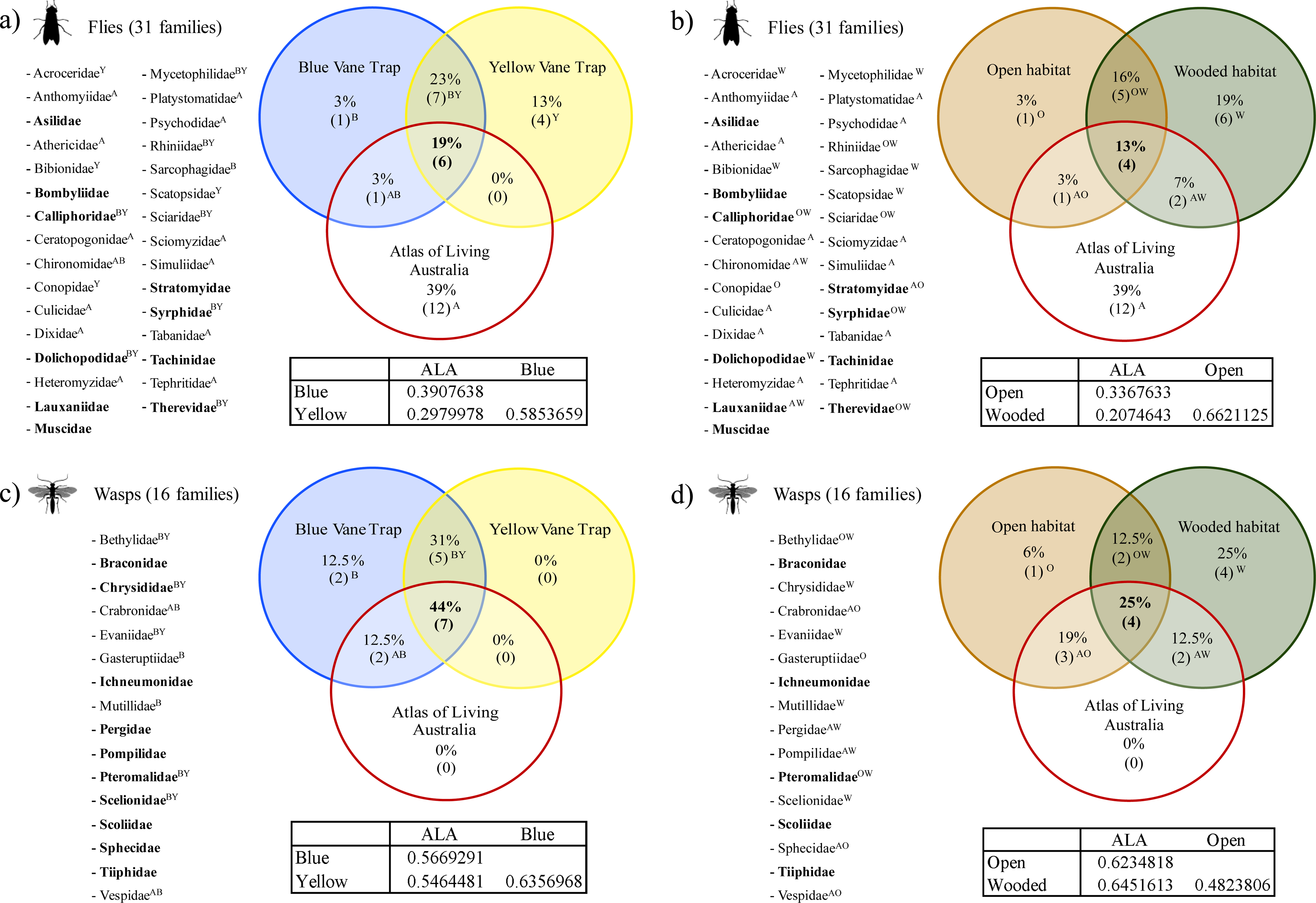
Comparison of the number and percentage of fly and wasp families sampled in our study using two different coloured traps - blue vane trap (BVT) and yellow vane trap (YVT) and within two different levels of wooded cover (habitat type) - Open and wooded, compared with those recorded on the Atlas of Living Australia (ALA) database for the sampling area (~3, 600 km^2^). Records shown for fly families by (a) trap colour and (b) habitat type, and for wasps by (c) trap colour and (d) habitat type. Letters beside names indicate where families were sampled - B=blue, Y=yellow, A=ALA, O=open, W=wooded. Tables below circles provide upper quantile values for the level of overlap between the ALA, BVTs and YVTs, calculated using the Morista-Horn index. A value of zero indicates no overlap, whilst a value of 1 indicates complete overlap.

BVTs had a higher probability of sampling wasp families compared with all available records on the ALA for the sampling area (Est = 3.26, SE = 1.56, P<0.01), however there was no difference in the likelihood for other variables tested (YVTs: Est = 0.78, SE = 0.76, P=0.28; Open: Est = 0.26, SE = 0.72, P=0.72; Wooded: Est = 0.85, SE = 0.77, P=0.27, Figs. 1c, 1d). Every family of wasp reported on the ALA (n=9) was also caught in BVTs, with seven additional families sampled using this colour that were not recorded on the ALA (Fig. 1c). Of the 16 families of wasps sampled, YVTs caught 12 families (Fig. 1c). Ten wasp families were sampled within open habitat types, including one unique family (Gasteruptiidae), while twelve families were collected in wooded habitats, including four unique families (Chrysididae, Evaniidae, Mutillidae, Scelionidae, Fig. 1d).

At the regional level encompassing our sampling area (11,500 km^2^), we found 39 fly and 13 wasp families recorded on the ALA (Fig. S2). Twenty fly families reported for the entire region on the ALA were not caught in vane traps during the study. Overall, fewer fly families were sampled in our study in both BVTs (Est = −1.99, SE = 0.53, P<0.01) and YVTs (Est = −1.78, SE = 0.53, P<0.01), along with both open (Est = −2.45, SE = 0.55, P<0.01) and wooded (Est = −1.78, SE = 0.53, P<0.01) habitat types compared with the wider region encompassing various habitat types and altitudinal gradients (Fig. S2). There was no difference in the number of wasp families recorded at the regional level on the ALA and those sampled in this study (BVTs: Est = 2.15, SE = 1.60, P=0.09; YVTs: Est = −0.33, SE = 0.84, P=0.69; Open: Est = −0.95, SE = 0.82, P=0.24; Wooded: Est = −0.37, SE = 0.86, P=0.67, Fig. S2). Every family of wasp reported on the ALA (13 families) at this scale was also caught in BVTs, with an additional family (Mutillidae) found that was not recorded on the ALA. Both BVTs and YVTs also sampled Bethylidae and Evaniidae, which were not recorded on the ALA (Fig. S2).

The majority of fly families sampled using all methods showed a peak activity period matching our sampling period of October-November (Fig. S3a), however six families in particular were either too rarely sampled to determine trends or showed different peak activity periods (Anthomyidae, Mycetophylidae, Platystomatidae, Sarcophagidae, Scatopsidae and Sciaridae, Fig. S3a). Around half of all wasp families recorded showed a similar activity peak to our sampling period, while most others were more active during the summer months (December-February) and Bethylidae had only low numbers recorded on the ALA, all within the first six months of the year (Fig. S3b).

### Trap colour

BVTs sampled greater average richness of all wasp and fly families combined, and greater abundance of all individuals across the study (Table 1, Figs. 2a, 2d). When modelled separately, all fly families also had greater average richness and abundance of individuals in BVTs (Table 1, Figs. 2b, 2e). Wasp families were sampled equally in both colours (Table 1, Figs. 2c, 2f).

**Table 1:** Differences by trap colour and habitat type of the richness of all fly (Diptera) and wasp (Hymenoptera) families and abundance of individuals, as well as the richness and abundance of each order when modelled separately. Bold values indicate a significant response. Reference categories used were *blue* for colour and *open* for habitat.

|  |  | Estimate | Std. Error | z value | Pr(>\|z\|) |
| --- | --- | --- | --- | --- | --- |
| *Richness* | Colour | -0.22 | 0.06 | -3.38 | **<0.01** |
|  | Habitat | -0.01 | 0.07 | -0.20 | 0.84 |
| *Abundance* | Colour | -0.79 | 0.10 | -7.80 | **<0.01** |
|  | Habitat | -0.35 | 0.11 | -3.06 | **<0.01** |
| *Fly richness* | Colour | -0.34 | 0.08 | -4.29 | **<0.01** |
|  | Habitat | -0.10 | 0.08 | -1.28 | 0.20 |
| *Fly abundance* | Colour | -0.87 | 0.11 | -8.23 | **<0.01** |
|  | Habitat | -0.43 | 0.12 | -3.48 | **<0.01** |
| *Wasp richness* | Colour | -0.07 | 0.13 | -0.57 | 0.57 |
|  | Habitat | 0.13 | 0.14 | 0.92 | 0.36 |
| *Wasp abundance* | Colour | -0.27 | 0.18 | -1.52 | 0.13 |
|  | Habitat | -0.02 | 0.19 | -0.09 | 0.93 |

**Figure 2:**
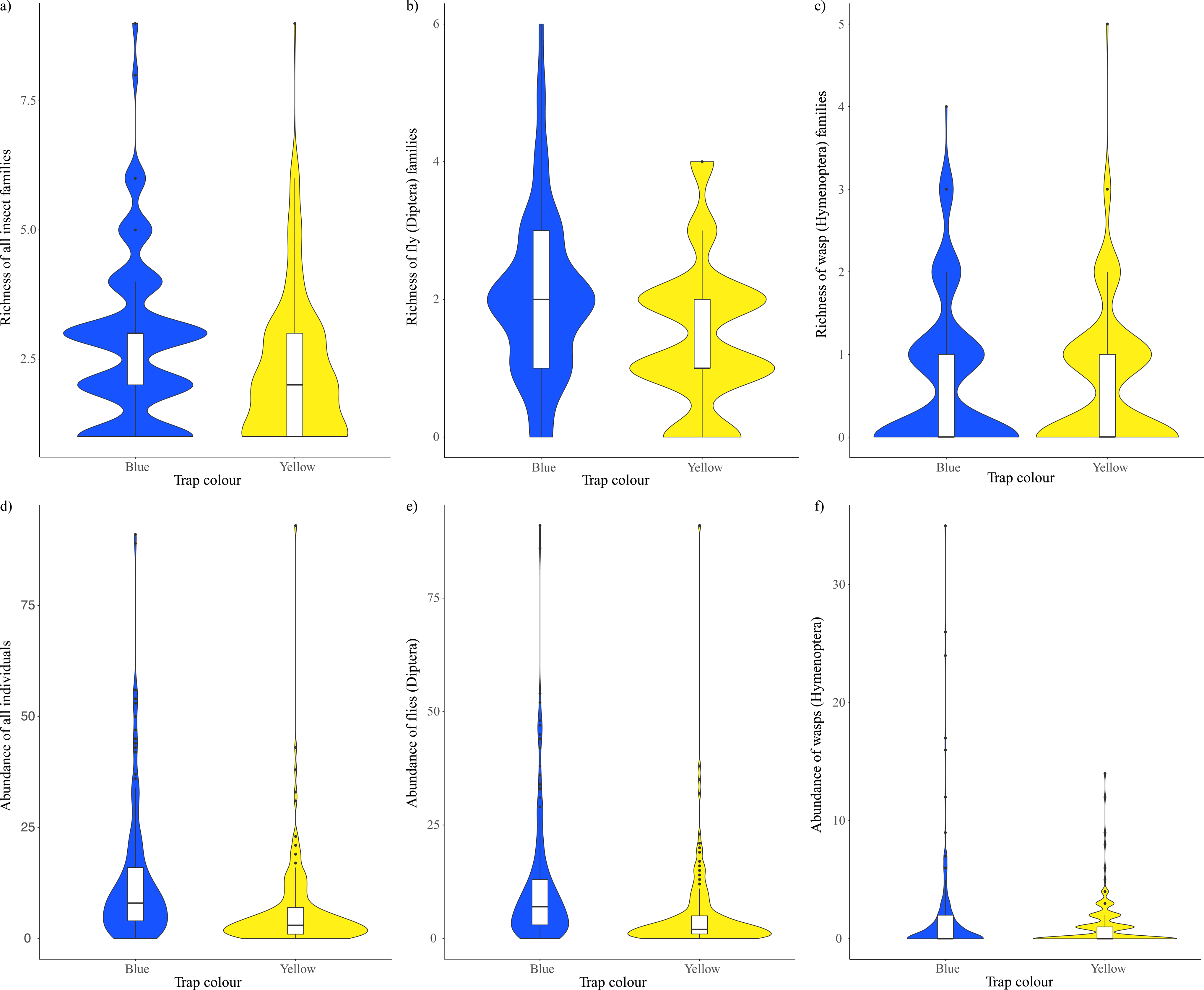
Average richness of (a) all insect families (flies and wasps), (b) only fly families and (c) only wasp families, trapped per site in blue versus yellow traps. Abundance of individuals for each of these groups shown in (d-f).

Of the nine families sampled with greater than 30 individuals, three were sampled more often in BVTs than YVTs (Diptera: Bombyliidae p<0.01, Syrphidae p<0.01, Hymenoptera: Scoliidae p<0.01; Table 2). Four families were sampled more often in YVTs (Diptera: Muscidae p<0.01, Tachinidae p=0.03, Therevidae p=0.02, Hymenoptera: Bethylidae p<0.01). There was no difference between colours for Braconidae or Tiphiidae (Hymenoptera) (Table 2).

**Table 2:**
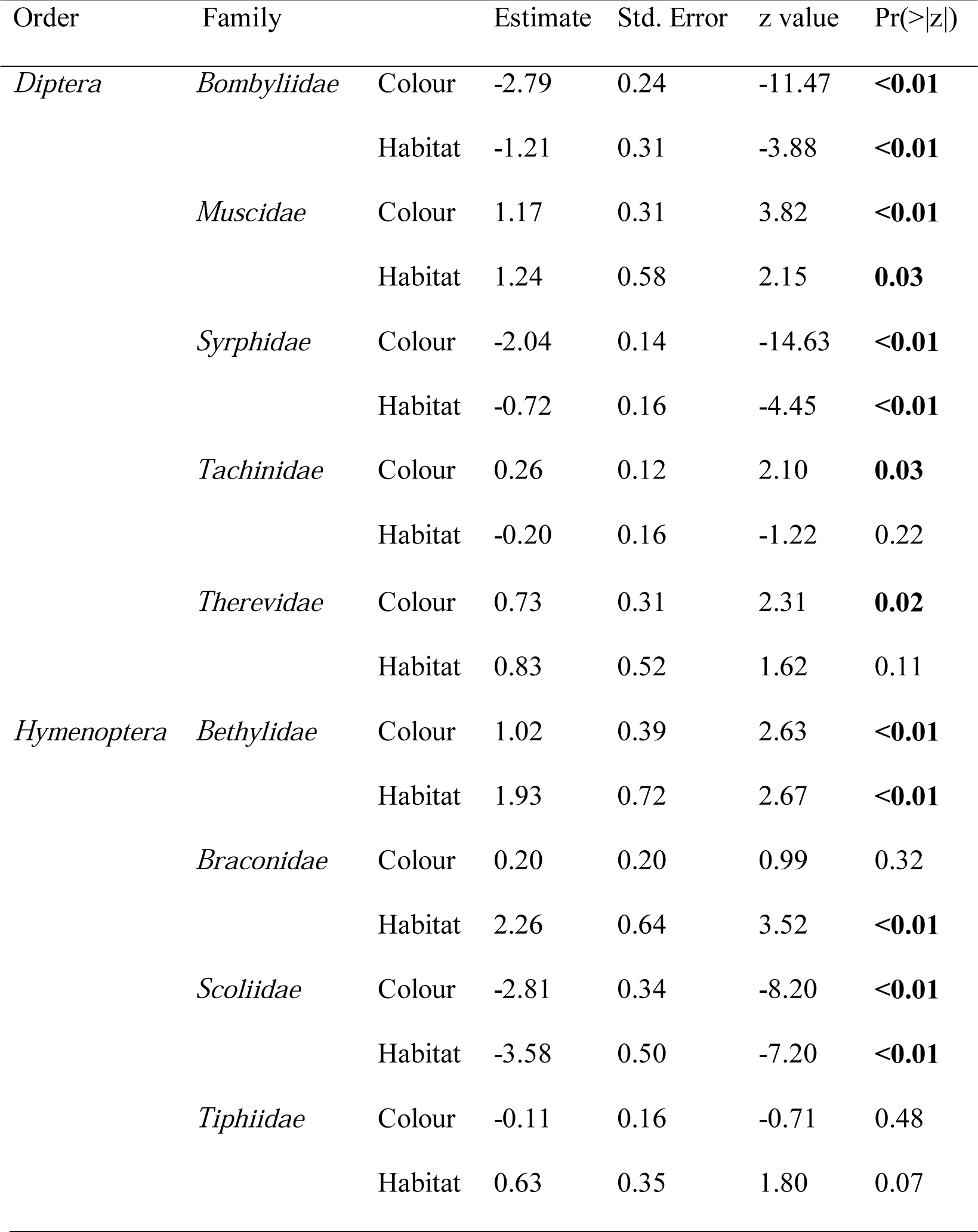
Differences by trap colour and habitat type of the abundance of individuals from fly (Diptera) and wasp (Hymenoptera) families sampled with >30 individuals. Bold values indicate a significant response. Reference categories used were *blue* for colour and *open* for habitat.

### Habitat

Average richness of all fly and wasp families did not differ between habitat type, even when modelled separately (Table 1, Figs. 3a-3c). However, the abundance of all flies and wasps was greater in open habitats (Table 1, Fig. 3d). This trend occurred when also modelling fly abundance separately (Table 1, Fig. 3e). Wasp abundance was sampled equally in both habitat types (Table 1, Fig. 3f).

**Figure 3:**
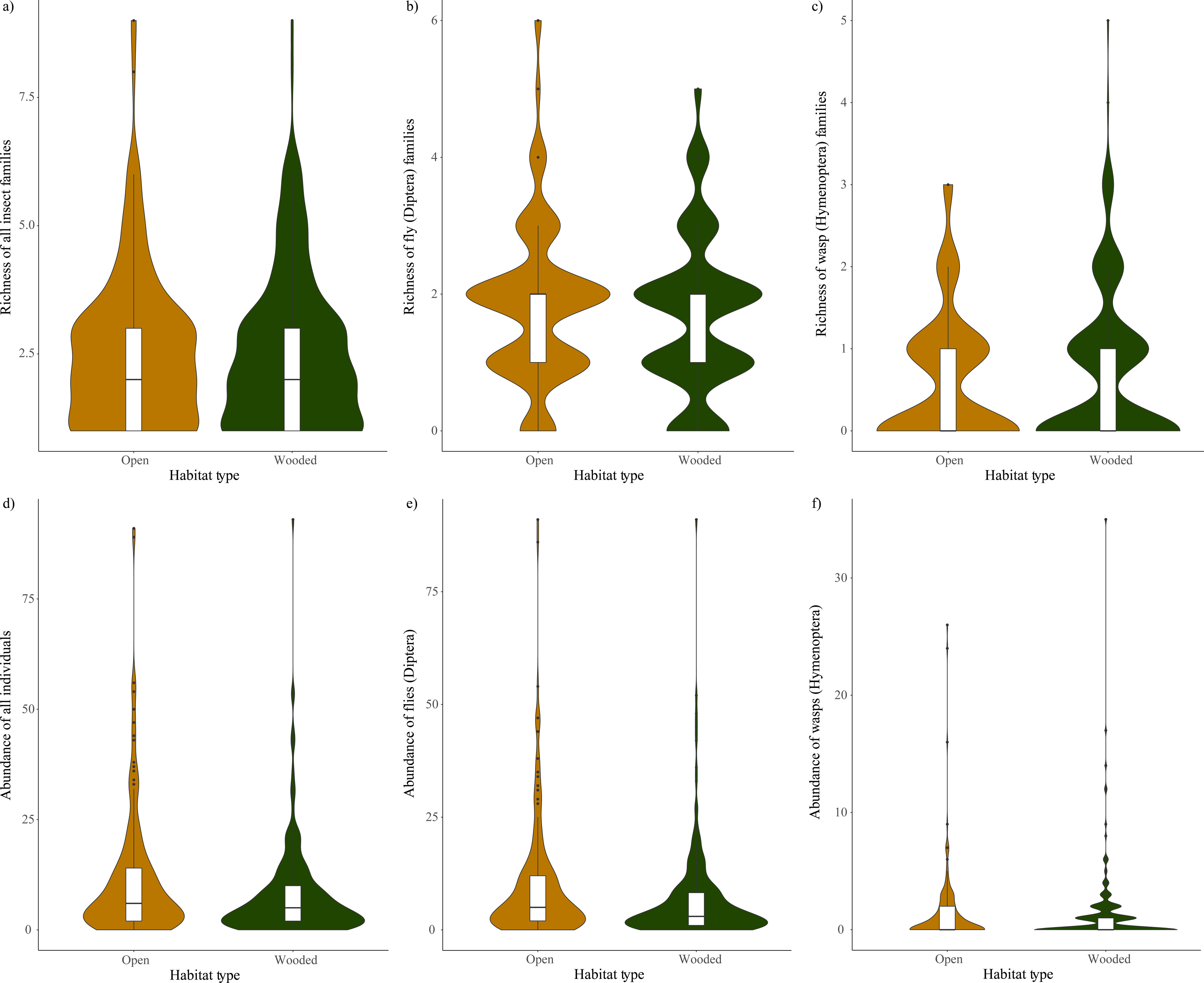
Average richness of (a) all insect families (flies and wasps), (b) only fly families and (c) only wasp families, trapped per site in open versus wooded habitats. Abundance of individuals for each of these groups shown in (d-f).

Of the nine families sampled with greater than 30 individuals, three families were found more often in open areas (Bombyliidae p<0.01, Syrphidae p<0.01 and Scoliidae p<0.01), whilst three families were more often sampled in wooded areas (Muscidae p=0.03, Bethylidae p<0.01, Braconidae p<0.01; Table 2). Three families showed no preference for either habitat type (Tachinidae, Therevidae, Tiphiidae; Table 2).

## DISCUSSION

This study demonstrates that vane traps can be used to representatively sample the non-bee pollinator fauna among different habitat types within agricultural landscapes. At the scale of the sampling area, we collected and identified an equal number of families within a short sampling period to that recorded on an historical database. When broadening the scale to encompass the region, we found our sampling to be equal for wasps, but less representative for the fly fauna, potentially due to the inclusion of larger forest patches, waterbodies and altitudinal gradients at this scale. Large numbers of individuals were collected from multiple fly and wasp families across the survey period, with certain families known to be pollinators (e.g. Syrphidae: Armstrong, 1979; Bombyliidae: Larson *et al.*, 2001; Scoliidae: Vithanage & Ironside, 1986) displaying a marked colour preference for BVTs. In contrast, family richness and abundance of insects was lower in YVTs, but this colour sampled some less common families important to the provision of multiple ecosystem services, including pollination (e.g. Acroceridae: Winterton, 2012; Bibionidae: D’Arcy-Burt & Blackshaw, 1991). These results, combined with similar findings for the bee fauna within this region (Hall, 2018), indicate that vane traps are an important survey tool to understand the demography and distribution of pollinator communities.

### Sampling effectiveness

Whilst many studies have compared different trapping methods and tested colour attraction in pollinating insects (Kirk, 1984; Campbell & Hanula, 2007; Saunders & Luck, 2013), few have directly compared results to the known insect fauna of the study region in question. Here, we not only compare two colours of vane traps (blue and yellow), but also measure the effectiveness of this sampling method compared to a baseline of insect diversity within the same study area. The ALA is a comprehensive repository of environmental and biological information for Australia (Atlas of Living Australia, 2019). It has been used to inform plant suitability under climate-change (Booth *et al.*, 2012; Belbin & Williams, 2016), but such databases are still rarely used as a reference point for insect communities within a geographic area (but see Godefroid *et al*., 2015; De Palma *et al*., 2017; Young *et al*., 2017 for examples of effective use of historical data obtained through databases and museum specimens). If we are to understand the distribution and dynamics of pollinator communities, surveys should be conducted across large spatial scales and compared with existing databases (Bartomeus *et al*., 2018). Vane traps appear well suited for such comparison studies.

At the scale of the sampling area, vane traps sampled an equivalent diversity of fly and wasp families to the ALA database. In fact, the diversity of wasp families captured was greater than that recorded for the study area. The number of fly families sampled was lower than, but comparable to, the ALA and sampled some families not recorded on it. The lower number of fly families sampled in our study is likely due to the short sampling period. We sampled during the austral spring which may not align with the peak activity of certain families, such as Anthomyiidae and Platystomatidae, which may more likely be recorded in early summer or autumn (Fig. S3a).

When expanded to the regional scale, the number of wasp families remained the same, however a number of additional fly families were recorded on the ALA that were not sampled in vane traps. There are a number of possible reasons for this. First, as discussed above, the main activity period of those additional fly families is different from our sampling period (e.g. Chloropidae, Drosophilidae), or they have been rarely recorded (e.g. only one record on the ALA for Pediciidae, Fig. S3a). Second, such families may be more often found within mountainous habitats, intact forest or waterbodies (e.g. Chloropidae, Drosophilidae, Empididae: Lambkin *et al*., 2011; Ephydridae: Keiper *et al*, 2002). These habitats were present within the greater regional area but not our sampling area, thus these families would be less likely to be found in more open, low altitude plains that comprise the less diverse habitats of our modified farm landscapes. Finally, families containing small-sized insects seem under-represented in our sampling. Such specimens were potentially missed when sorting traps due to their small size. However, these tiny insects are also likely to be missed in active surveys, such as sweep netting, and potentially unidentifiable when liquids or sticking agents are used with traps (Pickering & Stock, 2003).

### Colour preference

We found greater richness and abundance of non-bee pollinators in BVTs than in YVTs, indicating a strong colour preference to BVTs for most insect taxa sampled. This is consistent with other vane trap studies targeting bees (Stephen & Rao, 2005; Joshi *et al*., 2015; Hall, 2018) and is the first study comparing the effectiveness of these traps for sampling many of the non-bee fauna. Studies using other survey methods show some inconsistency in their ability to detect colour preference. For instance, Saunders & Luck (2013) found that yellow pan traps were preferred by pollinators (Hymenoptera and Diptera) across various habitats, but catches in each colour trap varied with habitat type. Similarly, Abrahamczyk *et al.* (2010) found that in tropical and subtropical forests, yellow traps sampled more non-formicid Hymenoptera than blue, mainly due to a few families—Crabronidae, Ichneumonidae, Nyssonidae and Pompilidae. However, Campbell & Hanula (2007) found that blue pan traps were more effective than yellow pan, standard Malaise or coloured Malaise traps in sampling flower visiting insects in forested ecosystems. Such inconsistencies may be attributable to differences in reflectance of colours under different environmental conditions.

The UV-reflectance of traps possibly influences colour preference by pollinating insects. Bees and hoverflies are known to respond to the UV reflectance of flowers (Koski & Ashman, 2014), and other vane trap studies have tested the influence of reflectance on colour choice by bees (Stephen & Rao, 2005; Joshi *et al*., 2015). Studies using other trapping methods have also tested UV reflectance, with varying results. For instance, Sircom *et al*. (2018) found different coloured pan traps used for bee surveys reflected to various degrees across wavelengths, with yellow traps generally having higher reflectance, while Shrestha *et al*. (2019) found hymenopterans showed no preference based on reflectance, but dipterans were generally more attracted to non-fluorescent than fluorescent traps. The preference for BVTs has been recorded in bees despite differing reports on the reflectance of each colour (Stephen & Rao, 2005; Joshi *et al*., 2015; Hall, 2018). In the present study, multiple non-bee families similarly preferred BVTs over YVTs, aligning with studies of bee communities. However, Koski & Ashman (2014) show that fluorescent patterning may be as important as colour, a feature present in many flowers but not on any common trapping method. We believe reflectance is an important quality in traps, and given the consistent colour of vane traps, they are well suited to sampling non-bee pollinators, particularly those that respond to UV reflectance. Incorporating fluorescent pattern into trap design could further improve their efficiency.

The ecological requirements of insects also likely affect attraction to different coloured traps (Kirk, 1984; Saunders & Luck, 2013). For instance, Syrphidae (hoverflies), which are known to mimic bees (particularly *Apis mellifera* L.) in their foraging behaviours (Knutson & Murphy, 1990; Golding & Edmunds, 2000, Campos-Jiménez *et al*., 2014), were sampled more often in blue-coloured traps compared to yellow by Campbell & Hanula (2007). This was supported by our finding of a seven-fold increase in abundance of this family in BVTs. In addition, Joshi *et al.* (2015) and Hall (2018) found that bees were mostly attracted by BVTs, suggesting that, having similar foraging behaviour to bees, syrphids may respond to the same visual cues (Kevan & Baker, 1983; Laubertie *et al*., 2006). Similarly, Bombyliidae (bee flies) are also known to forage on flowers (Kastinger & Weber, 2001; Larson *et al.*, 2001) and have some behavioural similarities to both syrphids and bees (Armstrong, 1979; Knutson & Murphy, 1990). Bombyliids also show greater preference for blue flowers over yellow (Kastinger & Weber, 2001) and have been more successfully trapped in blue pan traps than in yellow in forested ecosystems of south-eastern United States (Campbell & Hanula, 2007). One wasp family, Scoliidae (flower wasps), known to pollinate Macadamia and Orchids (Vithanage & Ironside 1986; Ciotek *et al.*, 2006), was also associated with BVTs in our study, presumably responding similarly to colour as the above taxa (Vuts *et al*., 2012).

In contrast, four families were found more often in YVTs. Muscidae and Tachinidae are quite large, generalist groups of flies, containing numerous species with different feeding requirements, including predatory, blood-sucking, parasitic and anthophilous behaviours (Larson *et al*., 2001; Skevington & Dang, 2002). Despite their apparent attraction to flowers, especially nectar, very little is known about their interactions with flowers (Larson *et al*., 2001). Therevidae contains numerous predatory flies (especially of coleopteran larvae), which mainly avoid flowers, although a few species appear to feed on nectar and insect secretions (Irwin & Lyneborg, 1989; Irwin 2001; Skevington & Dang, 2002; van Herk *et al*., 2015). Bethylidae are known as beneficial insects and parasitoids of Lepidoptera and Coleoptera (Danthanarayana, 1980; Berry, 1998). Smith *et al*. (2015) sampled them on yellow sticky traps, but other coloured traps were not used in this study. Little is known about any possible colour attraction for these two families.

Similarly, BVTs were particularly efficient at catching wasp families. The four families (Crabronidae, Gasteruptiidae, Mutillidae and Vespidae) only caught in BVTs comprise mostly predatory or parasitic species, but are also potential flower visitors, sometimes even foraging at night (Portman *et al.*, 2010; Wilson *et al.*, 2010). These may be missed when surveying only in daylight hours. A benefit of vane traps is that they can be left *in situ* night and day throughout the sampling period, maximizing the chances of sampling species with different foraging behaviours.

YVTs were effective at catching less common fly families, indicating both BVTs and YVTs might be required to comprehensively sample dipteran pollinators. Many of these species appear to require floral resources at least temporarily. For example, nectar collection by adult Chironomidae might enhance their mating success (Larson *et al.*, 2001; Høye *et al*., 2013). Many others are important pollinators as well as providing several additional ecosystem services, such as soil turnover, parasitism and scavenging (D’Arcy-Burt & Blackshaw, 1991; Larson *et al.*, 2001; Santos *et al*., 2008; Winterton, 2012). These services may be particularly important, or indeed drive the response of taxa, in different habitat types.

### Habitat association

Colour preference of pollinators has been compared in different habitat types across numerous studies. These include comparisons within different forested ecosystems (Campbell & Hanula, 2007; Abrahamczyk *et al*., 2010), native bush and crop (Saunders & Luck, 2013) or sunny versus shaded (vegetated) sites (Hoback *et al*., 1999). Such studies differ slightly from ours, in that wooded habitats here consisted of linear strips or scattered trees with an open understorey of coarse woody debris, while open habitats lacked trees and much of the woody litter.

In studies comparing open to closed habitats (forest, shade), microclimatic conditions, such as light, wind or humidity (Barradas & Fanjul, 1986), or reflectance and visibility of the coloured elements likely differ between habitat type (Abrahamczyk *et al.*, 2010). In contrast, due to the relatively sparse nature of wooded habitats within farmland, we believe differences between open and wooded habitats in our study are likely driven by provision of different resources (floral, nesting and prey) within each habitat type rather than microclimate or trap visibility.

Many studies stress the importance of taking into account habitat when choosing different trapping methods. Comparing coloured pan traps in native bush and almond orchard, Saunders & Luck (2013) highlighted the need to consider habitat due to no individual colour adequately trapping target insects across all habitats. Similarly, with sticky traps, Hoback *et al.* (1999) found diversity measures were not affected by trap colour but determined by position of traps (shaded or exposed). In the present study, family richness in open or wooded habitats did not differ, but abundance of non-bee pollinators was greater in open habitat, likely due to the three most abundant flower-visiting families preferring open areas.

For families present only in open habitats, several ecological traits could explain their presence. Some families (Diptera: Conopidae, Stratiomyidae; Hymenoptera: Crabronidae) are known flower visitors and would therefore be present where the flowering resources are most abundant (Conopidae: Armstrong, 1979; Stratiomyidae: Kevan & Baker, 1983; Crabronidae: Portman *et al.*, 2010). The three families that were associated most with BVTs (Bombyliidae, Syrphidae and Scoliidae) were also sampled more often in open habitats. Just as Hall *et al.* (2019) attributed the higher presence of bees in open habitats to greater floral resources, the presence of these families in open habitats could indicate behavioural similarities with bees.

Families more often sampled in wooded habitats were either associated with YVTs (Muscidae, Bethylidae) or displayed no colour preference (Braconidae). Overall, we found more unique families in wooded habitat which we attribute to a greater diversity of nest and perch sites, litter and food (e.g. insect and arachnid prey). For example, Lauxaniidae feed on leaf litter and are more likely found in the presence of shrubs, trees and leaves than in grasslands (Merz, 2004). Other families predate or parasitise caterpillars, orthopterans or spiders (Pompilidae: Rayor, 1996; Acroceridae: Winterton, 2012). Several families are known to parasitise solitary bee nests, wasp nests or cockroach ootheca (Sacrophagidae: Skevington & Dang, 2002; Evaniidae: Deyrup & Atkinson, 1993; Chrysididae: Rosenheim, 1987). Others make their nests on wood or fungi (Mycetophilidae: Madwar, 1937). These results further highlight the need to sample across multiple habitat types, as this might affect capture of certain taxa more than colour preference.

Vane traps are effective at sampling a diverse fauna present in habitats associated with modified agricultural landscapes. However, other habitat types such as wetlands, forests or mountains were present within the wider region and appear to greatly increase diversity of fly families in particular (Fig S2). It is therefore important to conduct sampling across various habitat types if wanting to ensure a representative insect census of an entire region. Due to consistency and quality of colour and ease of installation, vane traps seem an ideal method for comparisons between habitats.

### Conclusions and recommendations

As with any one trapping technique, there are pros and cons to using vane traps for sampling insect fauna. One positive is that due to placement height and the absence of detergents, vane traps used here were less likely to be disturbed, drunk, or spilled by animals (wild or cattle), which may be a hazard when using other methods, such as pan traps. However, vane traps have at times been suspected of oversampling the bee community (Gibbs *et al*., 2017). It is true that the number of individual bees captured in this region was relatively large for two months of sampling (Hall, 2018) and we had high abundance of certain families in this study, but not to the extent we would be concerned with oversampling. Regardless, caution is needed to limit the impact that trapping will have on the pollinator community. One possible solution would be to sample for a shorter period but to increase the number of sampling periods to capture seasonal changes (turnover) in the pollinator community (Thomsen *et al*., 2016; Winfree *et al*., 2018). This would further increase the effectiveness of this trapping method in sampling a representative pollinator insect community.

A further limitation is the general dearth of knowledge of insect pollinators, which makes it difficult to draw meaningful ecological conclusions, given even basic distributional data is often lacking for Australia. This highlights the importance of using historical datasets as a baseline comparison to determine how representative sampling is for an area (Bartomeus *et al*., 2018). Taxonomic keys to the region are also limiting in that they only go as far as family-level, and expertise on pollinators is hard to come by (Batley & Hogendoorn, 2009). Although identification at the species-level often allows for the greatest precision (Lenat & Resh, 2001), family-level identification provides better functional and behavioural indications than order, which is sometimes used for such studies. Indeed, clear trends appeared for three families of non-bee pollinators: Bombyliidae, Syrphidae (Diptera) and Scoliidae (Hymenoptera), which preferred BVTs here. However, it is difficult to establish colour preferences for more generalist families within Diptera, which may include species with very different feeding behaviours (e.g. flesh flies, flower flies, predatory flies). In such cases, it would be ideal to use species-level classification where sufficient data exists.

Vane traps are an effective sampling method to capture a representative non-bee pollinator community, including flies and wasps. Whereas bees displayed clear preference for BVTs, results of this study suggest that in order to sample the complete pollinator community, a range of colours may be more suitable. Additionally, to capture greater numbers of other flower visiting insects such as butterflies or beetles, it might be necessary to combine this with other trapping techniques such as Malaise traps. Active techniques, such as sweep netting could be employed if wanting to answer more specific questions about the function of communities, but for pollinator census, vane traps are a cheap and efficient method that can be used to good effect across multiple habitats.

## ACKNOWLEDGEMENTS

This work was conducted with financial assistance from the Holsworth Wildlife Research Endowment, awarded to MH during his PhD at La Trobe University. ER conducted later parts of this work as an intern at the Hawkesbury Institute for the Environment, under the supervision of James Cook. Many thanks to all the generous landholders for their interest and access to properties, volunteers for assistance in conducting insect surveys and to Andrew Bennett and Dale Nimmo for invaluable supervision and support (of MH) throughout the field component of this study.

## REFERENCES

Abrahamczyk, S., Steudel, B. & Kessler, M. (2010). Sampling Hymenoptera along a precipitation gradient in tropical forests: the effectiveness of different coloured pan traps. Entomologia Experimentalis et Applicata, 137, 262–268.

Alomar, D., González-Estévez, M.A., Traveset, A. & Lázaro, A. (2018). The intertwined effects of natural vegetation, local flower community, and pollinator diversity on the production of almond trees. Agriculture, Ecosystems and Environment, 264, 34–43.

Armstrong, J.A. (1979). Biotic pollination mechanisms in the Australian flora—a review. New Zealand Journal of Botany, 17, 467–508.

Atlas of Living Australia (ALA). (2019). Spatial portal. Accessed 22 January 2019. https://spatial.ala.org.au/

Barradas, V.L. & Fanjul, L. (1986). Microclimatic chacterization of shaded and open-grown coffee (Coffea arabica L.) plantations in Mexico. Agricultural and Forest Meteorology, 38, 101–112.

Bartholomew, C.S. & Prowell, D. (2005). Pan compared to malaise trapping for bees (Hymenoptera: Apoidea) in a longleaf pine savanna. Journal of the Kansas Entomological Society, 78, 390–392.

Bartomeus, I., Stavert, J.R., Ward, D. & Aguado, O. (2018). Historical collections as a tool for assessing the global pollination crisis. Philosophical Transactions of the Royal Society B, 374, 20170389.

Batley, M. & Hogendoorn, K. (2009). Diversity and conservation status of native Australian bees. Apidologie, 40, 347–354.

Belbin, L. & Williams, K.J. (2016). Towards a national bio-environmental data facility: experiences from the Atlas of Living Australia. International Journal of Geographical Information Science, 30, 108–125.

Bell, L.W. & Moore, A.D. (2012) Integrated crop–livestock systems in Australian agriculture: trends, drivers and implications. Agricultural Systems, 111, 1–12.

Berry, J.A. (1998). The bethyline species (Hymenoptera: Bethylidae: Bethylinae) imported into New Zealand for biological control of pest leafrollers. New Zealand Journal of Zoology, 25, 329–333.

Booth, T.H., Williams, K.J. & Belbin, L. (2012). Developing biodiverse plantings suitable for changing climatic conditions 2: Using the Atlas of Living Australia. Ecological Management & Restoration, 13, 274–281.

Brittain, C., Williams, N., Kremen, C. & Klein, A.M. (2013). Synergistic effects of non-Apis bees and honey bees for pollination services. Proceedings of the Royal Society of London B: Biological Sciences, 280, 20122767.

Bureau of Meteorology (2016). Climate data online. Accessed June 2016. http://www.bom.gov.au/climate/data/

Campbell, J.W. & Hanula, J.L. (2007) Efficiency of malaise traps and colored pan traps for collecting flower visiting insects from three forested ecosystems. Journal of Insect Conservation, 11, 399–408.

Campbell, J.W., Irvin, A., Irvin, H., Stanley-Stahr, C. & Ellis, J.D. (2016). Insect visitors to flowering buckwheat, Fagopyrum esculentum (Polygonales: Polygonaceae), in north-central Florida. Florida Entomologist, 99, 264–268.

Campos-Jiménez, J., Martínez, A.J., Golubov, J., García-Franco, J. & Ruiz-Montiel, C. (2014). Foraging behavior of Apis mellifera (Hymenoptera: Apidae) and Lycastrirhyncha nitens (Diptera: Syrphidae) on Pontederia sagittata (Commelinales: Pontederiaceae) on a disturbed site. Florida Entomologist, 97, 217–223.

Ciotek, L., Giorgis, P., Benitez-Vieyra, S., & Cocucci, A. A. (2006). First confirmed case of pseudocopulation in terrestrial orchids of South America: pollination of Geoblasta pennicillata (Orchidaceae) by Campsomeris bistrimacula (Hymenoptera, Scoliidae). Flora, 201, 365–369.

CSIRO (2018). Australian Insect Families: Hymenoptera, viewed 24 November 2018, http://anic.ento.csiro.au/insectfamilies/key.aspx?OrderID=27447andPageID=identifyandKeyID=27.

D’Arcy-Burt, S. & Blackshaw, R.P. (1991). Bibionids (Diptera: Bibionidae) in agricultural land: a review of damage, benefits, natural enemies and control. Annals of Applied Biology, 118, 695–708.

Danthanarayana, W. (1980). Parasitism of the light brown apple moth, Epiphyas Postvittana (Walker), by its larval ectoparasite, Goniozus Jacintae Farrugia (Hymenoptera: Bethylidae), in natural populations in Victoria. Australian Journal of Zoology, 28, 685–692.

De Palma, A., Kuhlmann, M., Bugter, R., Ferrier, S., Hoskins, A.J., Potts, S.G., Roberts, S.P., Schweiger, O. & Purvis, A. (2017). Dimensions of biodiversity loss: spatial mismatch in land□use impacts on species, functional and phylogenetic diversity of European bees. Diversity and Distributions, 23, 1435–1446.

Desouhant, E., Navel, S., Foubert, E., Fischbein, D., Théry, M. & Bernstein, C. (2010). What matters in the associative learning of visual cues in foraging parasitoid wasps: colour or brightness?. Animal Cognition, 13, 535–543.

Deyrup, M., & Atkinson, T. H. (1993). Survey of evaniid wasps (Hymenoptera: Evaniidae) and their cockroach hosts (Blattodea) in a natural Florida habitat. Florida Entomologist, 76, 589–593.

Environmental Conservation Council (2001). Box-Ironbark forests and woodlands investigation: final report. Environmental Conservation Council, East Melbourne, Victoria.

Gibbons, P., Lindenmayer, D.B., Fischer, J., Manning, A.D., Weinberg, A., Seddon, J., Ryan, P. & Barrett, G. (2008). The future of scattered trees in agricultural landscapes. Conservation Biology, 22, 1309–1319.

Gibbs, J., Joshi, N.K., Wilson, J.K., Rothwell, N.L., Powers, K., Haas, M., Gut, L., Biddinger, D.J. & Isaacs, R. (2017). Does passive sampling accurately reflect the bee (Apoidea: Anthophila) communities pollinating apple and sour cherry orchards?. Environmental Entomology, 46, 579–588.

Godefroid, M., Cruaud, A., Rossi, J.P. & Rasplus, J.Y. (2015). Assessing the risk of invasion by Tephritid fruit flies: intraspecific divergence matters. PloS one, 10, e0135209.

Golding, Y.C. & Edmunds, M. (2000). Behavioural mimicry of honeybees (Apis mellifera) by droneflies (Diptera: Syrphidae: Eristalis spp.). Proceedings of the Royal Society of London B: Biological Sciences, 267, 903–909.

Hall, M. (2018). Blue and yellow vane traps differ in their sampling effectiveness for wild bees in both open and wooded habitats. Agricultural and Forest Entomology, 20, 487–495.

Hall, M.A., Nimmo, D.G., Cunningham, S.A., Walker, K. & Bennett, A.F. (2019). The response of wild bees to tree cover and rural land use is mediated by species’ traits. Biological Conservation, 231, 1–12.

Hall, M., Nimmo, D., Watson, S. & Bennett, A.F. (2018). Linear habitats in rural landscapes have complementary roles in bird conservation. Biodiversity and Conservation, 27, 2605–2623.

Heard, T.A. (1999). The role of stingless bees in crop pollination. Annual Review of Entomology, 44, 183–206.

Heinze, G. & Schemper, M. (2002). A solution to the problem of separation in logistic regression. Statistics in Medicine, 21, 2409–2419.

Heinze, G., Ploner, M., Dunkler, D. & Southworth, H. (2018). logistf: Firth’s bias reduced logistic regression. R package version, 1.23. https://rdrr.io/cran/logistf/man/logistf.html

Hoback, W.W., Svatos, T.M., Spomer, S.M. & Higley, L.G. (1999). Trap color and placement affects estimates of insect family-level abundance and diversity in a Nebraska salt marsh. Entomologia Experimentalis et Applicata, 91, 393–402.

Høye, T.T., Post, E., Schmidt, N.M., Trøjelsgaard, K. & Forchhammer, M.C. (2013). Shorter flowering seasons and declining abundance of flower visitors in a warmer Arctic. Nature Climate Change, 3, 759–763.

Inouye, D.W. & Pyke, G.H. (1988). Pollination biology in the Snowy Mountains of Australia: comparisons with montane Colorado, USA. Australian Journal of Ecology, 13, 191–205.

Irwin, M.E. (2001). Species composition and seasonal flight periodicity of stiletto flies (Diptera: Therevidae) occurring along the Kuiseb River, Gobabeb, Namibia. Cimbebasia, 17, 169–175.

Irwin, M.E. & Lyneborg, L. (1989). Family Therevidae. In Catalog of Diptera of the Australasian and Oceanian Regions. (Ed. NL Evenhuis.) pp. 353–358.

Joshi, N.K., Leslie, T., Rajotte, E.G., Kammerer, M.A., Otieno, M. & Biddinger, D.J. (2015). Comparative trapping efficiency to characterize bee abundance, diversity, and community composition in apple orchards. Annals of the Entomological Society of America, 108, 785–799.

Kastinger, C. & Weber, A. (2001). Bee-flies (Bombylius spp., Bombyliidae, Diptera) and the pollination of flowers. Flora, 196, 3–25.

Keiper, J. B., Walton, W. E., & Foote, B. A. (2002). Biology and ecology of higher Diptera from freshwater wetlands. Annual Review of Entomology, 47, 207–232.

Kennedy, C.M., Lonsdorf, E., Neel, M.C., Williams, N.M., Ricketts, T.H., Winfree, R., Bommarco, R., Brittain, C., Burley, A.L., Cariveau, D. & Carvalheiro, L.G. (2013). A global quantitative synthesis of local and landscape effects on wild bee pollinators in agroecosystems. Ecology Letters, 16, 584–599.

Kevan, P.G. & Baker, H.G. (1983). Insects as flower visitors and pollinators. Annual Review of Entomology, 28, 407–453.

Kimoto, C., DeBano, S.J., Thorp, R.W., Rao, S. & Stephen, W.P. (2012). Investigating temporal patterns of a native bee community in a remnant North American bunchgrass prairie using blue vane traps. Journal of Insect Science, 12, 1–23.

Kirk, W.D. (1984). Ecologically selective coloured traps. Ecological Entomology, 9, 35–41.

Kitching, R.L., Li, D. & Stork, N.E. (2001). Assessing biodiversity ‘sampling packages’: how similar are arthropod assemblages in different tropical rainforests?. Biodiversity and Conservation, 10, 793–813.

Klein, A.M., Vaissiere, B.E., Cane, J.H., Steffan-Dewenter, I., Cunningham, S.A., Kremen, C. & Tscharntke, T. (2006). Importance of pollinators in changing landscapes for world crops. Proceedings of the Royal Society B: Biological Sciences, 274, 303–313.

Knutson, L.V. & Murphy, W.L. (1990). Chapter 8: Insects: Diptera (Flies). In Morse, R.A. and Nowogrodzki, R. (eds). Honey bee pests, predators, and diseases (2^nd^ Ed). Cornell University Press.

Koh, I., Lonsdorf, E.V., Williams, N.M., Brittain, C., Isaacs, R., Gibbs, J. & Ricketts, T.H. (2016). Modeling the status, trends, and impacts of wild bee abundance in the United States. Proceedings of the National Academy of Sciences, 113, 140–145.

Koski, M.H. & Ashman, T.L. (2014). Dissecting pollinator responses to a ubiquitous ultraviolet floral pattern in the wild. Functional Ecology, 28, 868–877.

Kremen, C., Williams, N.M. & Thorp, R.W. (2002). Crop pollination from native bees at risk from agricultural intensification. Proceedings of the National Academy of Sciences, 99, 16812–16816.

Lambkin, C.L., Boulter, S.L., Starick, N.T., Cantrell, B.K., Bickel, D.J., Wright, S.G., Power, N., Schutze, M.K., Turco, F., Nakamura, A. & Burwell, C.J. (2011). Altitudinal and seasonal variation in the family-level assemblages of flies (Diptera) in an Australian subtropical rainforest: one hundred thousand and counting!. Memoirs of the Queensland Museum, 55, 315–331.

Larson, B. M. H., Kevan, P. G., & Inouye, D. W. (2001). Flies and flowers: taxonomic diversity of anthophiles and pollinators. The Canadian Entomologist, 133, 439–465.

Larsson, M. & Franzen, M. (2008). Estimating the population size of specialised solitary bees. Ecological Entomology, 33, 232–238.

Laubertie, E.A., Wratten, S.D. & Sedcole, J.R. (2006). The role of odour and visual cues in the pan□trap catching of hoverflies (Diptera: Syrphidae). Annals of Applied Biology, 148, 173–178.

Lefebvre, V., Villemant, C., Fontaine, C. & Daugeron, C. (2018). Altitudinal, temporal and trophic partitioning of flower-visitors in Alpine communities. Scientific Reports, 8, 4706.

Lenat, D.R. & Resh, V.H. (2001). Taxonomy and stream ecology—the benefits of genus-and species-level identifications. Journal of the North American Benthological Society, 20, 287–298.

Lentini, P.E., Martin, T.G., Gibbons, P., Fischer, J. & Cunningham, S.A. (2012) Supporting wild pollinators in a temperate agricultural landscape: maintaining mosaics of natural features and production. Biological Conservation, 149, 84–92.

McCravy, K. (2018). A review of sampling and monitoring methods for beneficial arthropods in agroecosystems. Insects, 9, 170.

McHugh, J.V. & Liebherr, J.K., (2009). Coleoptera:(beetles, weevils, fireflies). In Encyclopedia of Insects (2^nd^ Ed), pp.183–201.

Madwar, S. (1937). I-Biology and morphology of the immature stages of Mycetophilidae (Diptera, Nematocera). Philosophical Transactions of the Royal Society of London. Series B, Biological Sciences, 227, 1–110.

Marshall, S.A. (2017). Key to Diptera Families - Adults. In Kirk-Spriggs, A.H. and Sinclair, B.J. Manual of Afrotropical Diptera. Volume 1. Introductory chapters and keys to Diptera families. Suricata 4. South African National Biodiversity Institute, Pretoria, pp. 267–356.

Merz, B. (2004). Revision of the Minettia fasciata species-group (Diptera, Lauxaniidae). Revue Suisse de Zoologie, 111, 183–211.

Missa, O., Basset, Y., Alonso, A., Miller, S.E., Curletti, G., De Meyer, M., Eardley, C., Mansell, M.W. & Wagner, T. (2009). Monitoring arthropods in a tropical landscape: relative effects of sampling methods and habitat types on trap catches. Journal of Insect Conservation, 13, 103.

Morse, R.A. & Calderone, N.W. (2000). The value of honey bees as pollinators of US crops in 2000. Bee Culture, 128, 1–15.

Ollerton, J. (2017). Pollinator diversity: distribution, ecological function, and conservation. Annual Review of Ecology, Evolution, and Systematics, 48, 353–376.

Pickering, C.M. & Stock, M. (2003). Insect colour preference compared to flower colours in the Australian Alps. Nordic Journal of Botany, 23, 217–223.

Pietrzak, M., Seweryn, M. & Rempala, G. (2016). divo: tools for analysis of diversity and similarity in biological systems. R package version 0.1.2, http://CRAN.R-project.org/package=divo.

Pinheiro, J., Bates, D., DebRoy, S., Sarkar, D. & R Core Team (2018). nlme: linear and nonlinear mixed effects models. R package version 3.1-137, https://CRAN.R-project.org/package=nlme.

Portman, S.L., Frank, J. H., McSorley, R., & Leppla, N. C. (2010). Nectar-seeking and host-seeking by Larra bicolor (Hymenoptera: Crabronidae), a parasitoid of Scapteriscus mole crickets (Orthoptera: Gryllotalpidae). Environmental Entomology, 39, 939–943.

Potts, S.G., Imperatriz-Fonseca, V., Ngo, H.T., Biesmeijer, J.C., Breeze, T.D., Dicks, L.V., Garibaldi, L.A., Hill, R., Settele, J. & Vanbergen, A.J. (2016). The assessment report on pollinators, pollination and food production: summary for policymakers. Secretariat of the Intergovernmental Science-Policy Platform on Biodiversity and Ecosystem Services, Bonn, Germany..

R Core Team (2018). R: A Language and Environment for Statistical Computing. R Foundation for Statistical Computing, Vienna, Austria.

Rader, R., Bartomeus, I., Garibaldi, L.A., Garratt, M.P., Howlett, B.G., Winfree, R., Cunningham, S.A., Mayfield, M.M., Arthur, A.D., Andersson, G.K. & Bommarco, R. (2016). Non-bee insects are important contributors to global crop pollination. Proceedings of the National Academy of Sciences, 113, 146–151.

Rayor, L.S. (1996). Attack strategies of predatory wasps (Hymenoptera: Pompilidae; Sphecidae) on colonial orb web-building spiders (Araneidae: Metepeira incrassata). Journal of the Kansas Entomological Society, 69, 67–75.

Rosenheim, J.A. (1987). Host location and exploitation by the cleptoparasitic wasp Argochrysis armilla: the role of learning (Hymenoptera: Chrysididae). Behavioral Ecology and Sociobiology, 21, 401–406.

Santos, A.M., Serrano, J.C., Couto, R.M., Rocha, L.S., Mello-Patiu, C.A. & Garófalo, C.A. (2008). Conopid flies (Diptera: Conopidae) parasitizing Centris (Heterocentris) analis (Fabricius) (Hymenoptera: Apidae, Centridini). Neotropical Entomology, 37, 606–608.

Saunders, M.E. & Luck, G.W. (2013) Pan trap catches of pollinator insects vary with habitat. Australian Journal of Entomology, 52, 106–113.

Shrestha, M., Garcia, J.E., Chua, J.H., Howard, S.R., Tscheulin, T., Dorin, A., Nielsen, A. & Dyer, A.G. (2019). Fluorescent pan traps affect the capture rate of insect orders in different ways. Insects, 10, 40.

Sircom, J., Jothi, G.A. & Pinksen, J. (2018). Monitoring bee populations: are eusocial bees attracted to different colours of pan trap than other bees?. Journal of Insect Conservation, 22, 433–441.

Skevington, J.H. & Dang, P.T. (2002). Exploring the diversity of flies (Diptera). Biodiversity, 3, 3–27.

Smith, I.M., Hoffmann, A.A. & Thomson, L.J. (2015). Ground cover and floral resources in shelterbelts increase the abundance of beneficial hymenopteran families. Agricultural and Forest Entomology, 17, 120–128.

Stavert, J.R., Pattemore, D.E., Bartomeus, I., Gaskett, A.C. & Beggs, J.R. (2018). Exotic flies maintain pollination services as native pollinators decline with agricultural expansion. Journal of Applied Ecology, 55, 1737–1746.

Stephen,W.P. & Rao, S. (2005) Unscented color traps for non-Apis bees (Hymenoptera: Apiformes). Journal of the Kansas Entomological Society, 78, 373–380.

Stephen, W.P. & Rao, S. (2007) Sampling native bees in proximity to a highly competitive food resource (Hymenoptera: Apiformes). Journal of the Kansas Entomological Society, 80, 369–376.

Sullivan, R. (1981). Insect swarming and mating. The Florida Entomologist, 64, 44–65.

Taki, H., Murao, R., Mitai, K. & Yamaura, Y. (2018). The species richness/abundance–area relationship of bees in an early successional tree plantation. Basic and Applied Ecology, 26, 64–70.

Thomsen, P.F., Jørgensen, P.S., Bruun, H.H., Pedersen, J., Riis□Nielsen, T., Jonko, K., Słowińska, I., Rahbek, C. & Karsholt, O. (2016). Resource specialists lead local insect community turnover associated with temperature– analysis of an 18□year full□seasonal record of moths and beetles. Journal of Animal Ecology, 85, 251–261.

Thomson, D.M. (2019). Effects of long-term variation in pollinator abundance and diversity on reproduction of a generalist plant. Journal of Ecology, 107, 491–502.

van Herk, W.G., Vernon, R.S., Cronin, E.M.L. & Gaimari, S.D. (2015). Predation of Thereva nobilitata (Fabricius) (Diptera: Therevidae) on Agriotes obscurus L. (Coleoptera: Elateridae). Journal of Applied Entomology, 139, 154–157.

Venables, W.N. & Ripley, B.D. (2002) Modern applied statistics with S. (4th Ed). Springer, New York.

Vithanage, V., & Ironside, D. A. (1986). The insect pollinators of macadamia and their relative importance. Journal of the Australian Institute of Agricultural Science, 52, 155–160.

Vrdoljak, S.M. & Samways, M.J. (2012). Optimising coloured pan traps to survey flower visiting insects. Journal of Insect Conservation, 16, 345–354.

Vuts, J., Razov, J., Kaydan, M.B. & Tóth, M. (2012). Visual and olfactory cues for catching parasitic wasps (Hymenoptera: Scoliidae). Acta Zoologica Academiae Scientiarum Hungaricae, 58, 351–359.

Westphal, C., Bommarco, R., Carré, G., Lamborn, E., Morison, N., Petanidou, T., Potts, S.G., Roberts, S.P., Szentgyörgyi, H., Tscheulin, T. & Vaissière, B.E. (2008). Measuring bee diversity in different European habitats and biogeographical regions. Ecological Monographs, 78, 653–671.

Wilson, J.S., Williams, K.A., Tanner, D.A. & Pitts, J.P. (2010). Nectaring by nocturnal velvet ants (Hymenoptera: Mutillidae). The Southwestern Naturalist, 55, 441–443.

Winfree, R., Reilly, J.R., Bartomeus, I., Cariveau, D.P., Williams, N.M. & Gibbs, J. (2018). Species turnover promotes the importance of bee diversity for crop pollination at regional scales. Science, 359, 791–793.

Winterton, S.L. (2012). Review of Australasian spider flies (Diptera, Acroceridae) with a revision of Panops Lamarck. ZooKeys, 172, 7–75.

Young, B.E., Auer, S., Ormes, M., Rapacciuolo, G., Schweitzer, D. & Sears, N. (2017). Are pollinating hawk moths declining in the Northeastern United States? An analysis of collection records. PloS one, 12, e0185683.

Zhang, W., Ricketts, T.H., Kremen, C., Carney, K. & Swinton, S.M. (2007). Ecosystem services and dis-services to agriculture. Ecological Economics, 64, 253–260.

Zuur, A.F., Hilbe, J.M. & Ieno, E.N. (2013). A beginner’s Guide to GLM and GLMM with R. Highland Statistics Limited, U.K.

