## Supplementary material for "High sampling effectiveness for non-bee pollinators using vane traps in both open and wooded habitats"

**SUPPORTING INFORMATION**

Table S1: Abundance of all individuals from 19 fly (Diptera) and 16 wasp (Hymenoptera) families collected using blue and yellow vane traps in both open and wooded habitats. Families in bold had >30 individuals collected across the study.

|  |  | **Open** | | **Wooded** | |
| --- | --- | --- | --- | --- | --- |
| Order | Family | **Blue** | **Yellow** | **Blue** | **Yellow** |
| **Diptera** |  |  |  |  |  |
|  | Acroceridae |  |  |  | 1 |
|  | Asilidae |  | 2 | 11 | 14 |
|  | Bibionidae |  |  |  | 1 |
|  | **Bombyliidae** | 204 | 12 | 88 | 6 |
|  | Calliphoridae | 8 | 7 | 3 | 7 |
|  | Chironomidae |  |  | 1 |  |
|  | Conopidae |  | 1 |  |  |
|  | Dolichopodidae |  |  | 1 | 1 |
|  | Lauxaniidae |  |  | 1 | 2 |
|  | **Muscidae** | 3 | 12 | 12 | 32 |
|  | Mycetophilidae |  |  | 3 | 4 |
|  | Rhiniidae | 13 | 3 | 3 | 1 |
|  | Sarcophagidae |  |  | 1 |  |
|  | Scatopsidae |  |  |  | 2 |
|  | Sciaridae | 1 |  | 8 | 7 |
|  | Stratiomyidae | 1 | 1 |  |  |
|  | **Syrphidae** | 750 | 104 | 465 | 77 |
|  | **Tachinidae** | 277 | 285 | 259 | 373 |
|  | **Therevidae** | 3 | 7 | 15 | 21 |
| Abundance |  | **1260** | **434** | **871** | **549** |
| **Hymenoptera** |  |  |  |  |  |
|  | **Bethylidae** | 1 | 2 | 12 | 19 |
|  | **Braconidae** | 3 | 2 | 45 | 50 |
|  | Chrysididae |  |  | 6 | 1 |
|  | Crabronidae | 1 |  |  |  |
|  | Evaniidae |  |  | 1 | 1 |
|  | Gasteruptiidae | 1 |  |  |  |
|  | Ichneumonidae | 2 |  | 1 | 5 |
|  | Mutillidae |  |  | 2 |  |
|  | Pergidae |  |  | 1 | 1 |
|  | Pompilidae |  |  | 2 | 3 |
|  | Pteromalidae |  | 1 | 2 | 2 |
|  | Scelionidae |  |  | 2 | 2 |
|  | **Scoliidae** | 145 | 9 | 5 |  |
|  | Sphecidae | 10 | 19 |  |  |
|  | **Tiphiidae** | 20 | 19 | 66 | 58 |
|  | Vespidae | 6 |  |  |  |
| Abundance |  | **189** | **52** | **145** | **142** |
| **Total abundance** |  | **1449** | **486** | **1016** | **691** |

Table S2: Abundance of all individuals from the 5 fly (Diptera) and 4 wasp (Hymenoptera) families with >30 individuals collected. Also stated are the number and percentage of total traps these families were caught in.

| Order | Family | # individuals | # traps | % trap sampled in |
| --- | --- | --- | --- | --- |
| *Diptera* | *Bombyliidae* | 310 | 62 | 16% |
|  | *Muscidae* | 59 | 34 | 9% |
|  | *Syrphidae* | 1396 | 232 | 60% |
|  | *Tachinidae* | 1194 | 199 | 52% |
|  | *Therevidae* | 46 | 32 | 8% |
| *Hymenoptera* | *Bethylidae* | 34 | 27 | 7% |
|  | *Braconidae* | 100 | 41 | 11% |
|  | *Scoliidae* | 159 | 46 | 12% |
|  | *Tiphiidae* | 163 | 69 | 18% |

Figure S1: (a) The study region (black dots represent study landscapes and grey shading denotes native tree cover), set within Victoria, Australia. Blue and yellow vane traps were hung at a height of approximately two metres from (b) tree branches at wooded sites or (c) poles at open sites.

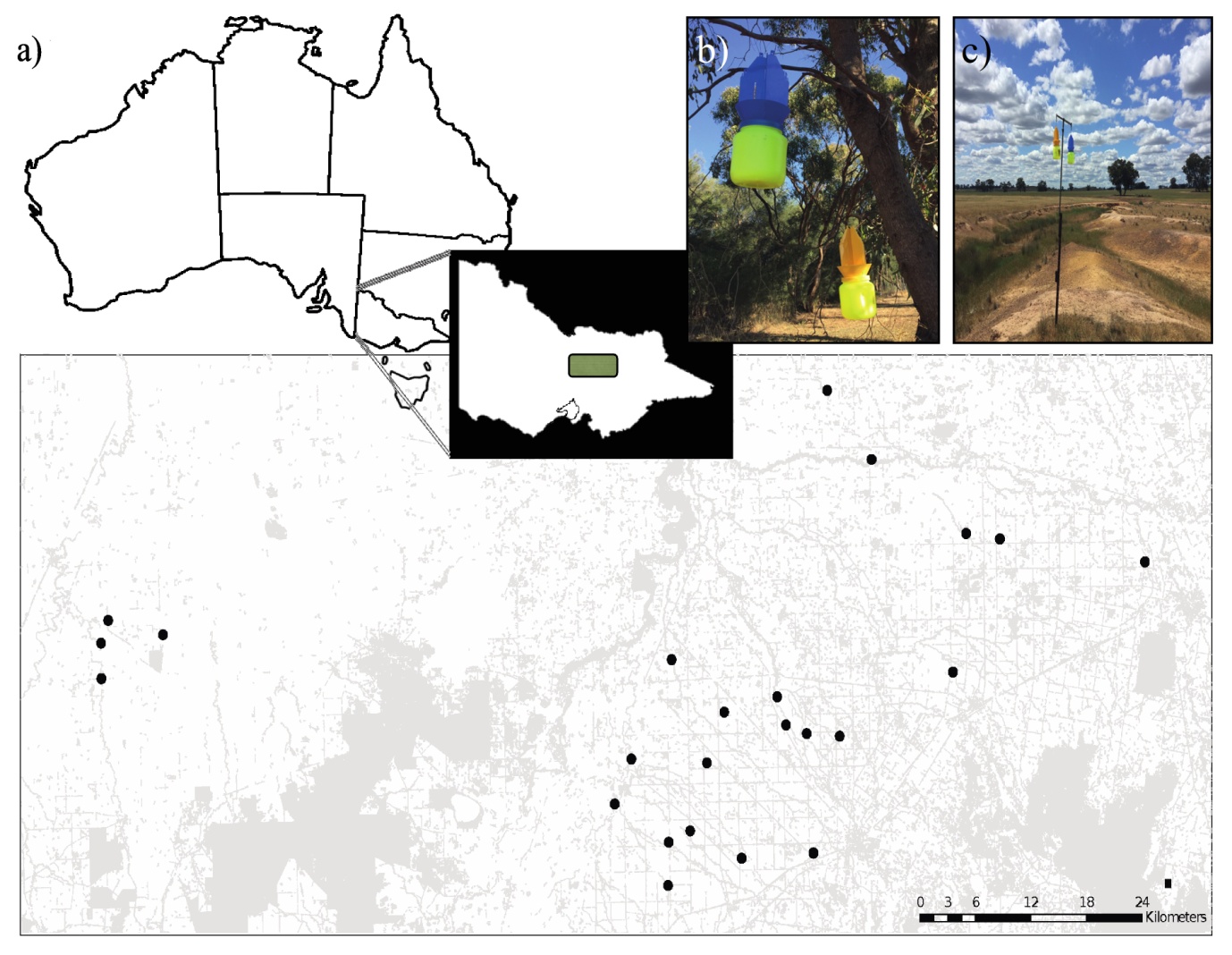

Figure S2: Comparison of the number of fly and wasp families sampled in our study, compared with those recorded on the Atlas of Living Australia (ALA) database for the entire region (11, 550 km^2^). Records shown for fly families by (a) trap colour and (b) habitat type, and for wasps by (c) trap colour and (d) habitat type. Codes indicate where families were sampled - B=blue, Y=yellow, A=ALA, O=open, W=wooded.

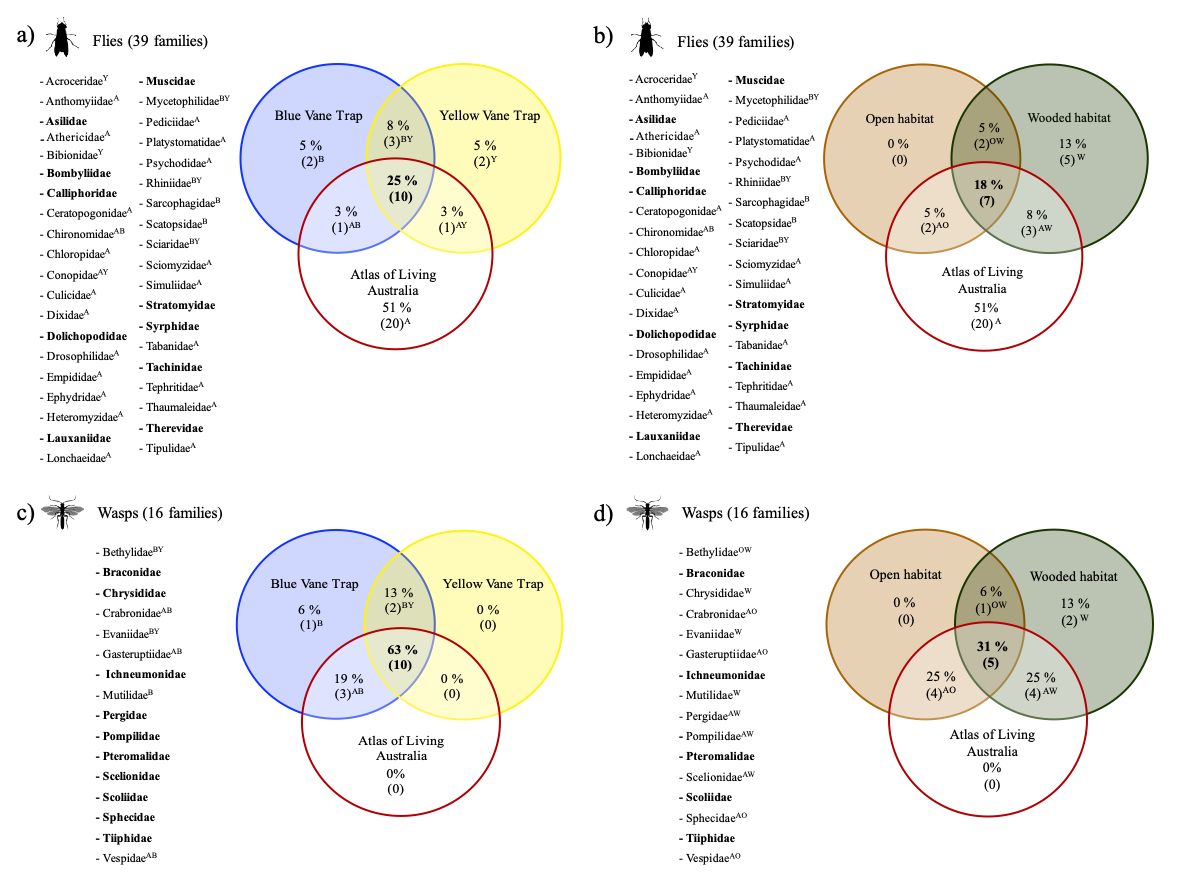

Figure S3: Number of records of each family of (a) fly and (b) wasp from the Atlas of Living Australia (ALA) database by month, used here as a proxy for peak foraging period. Records for the state of Victoria only are shown, covering all time periods.

(a)

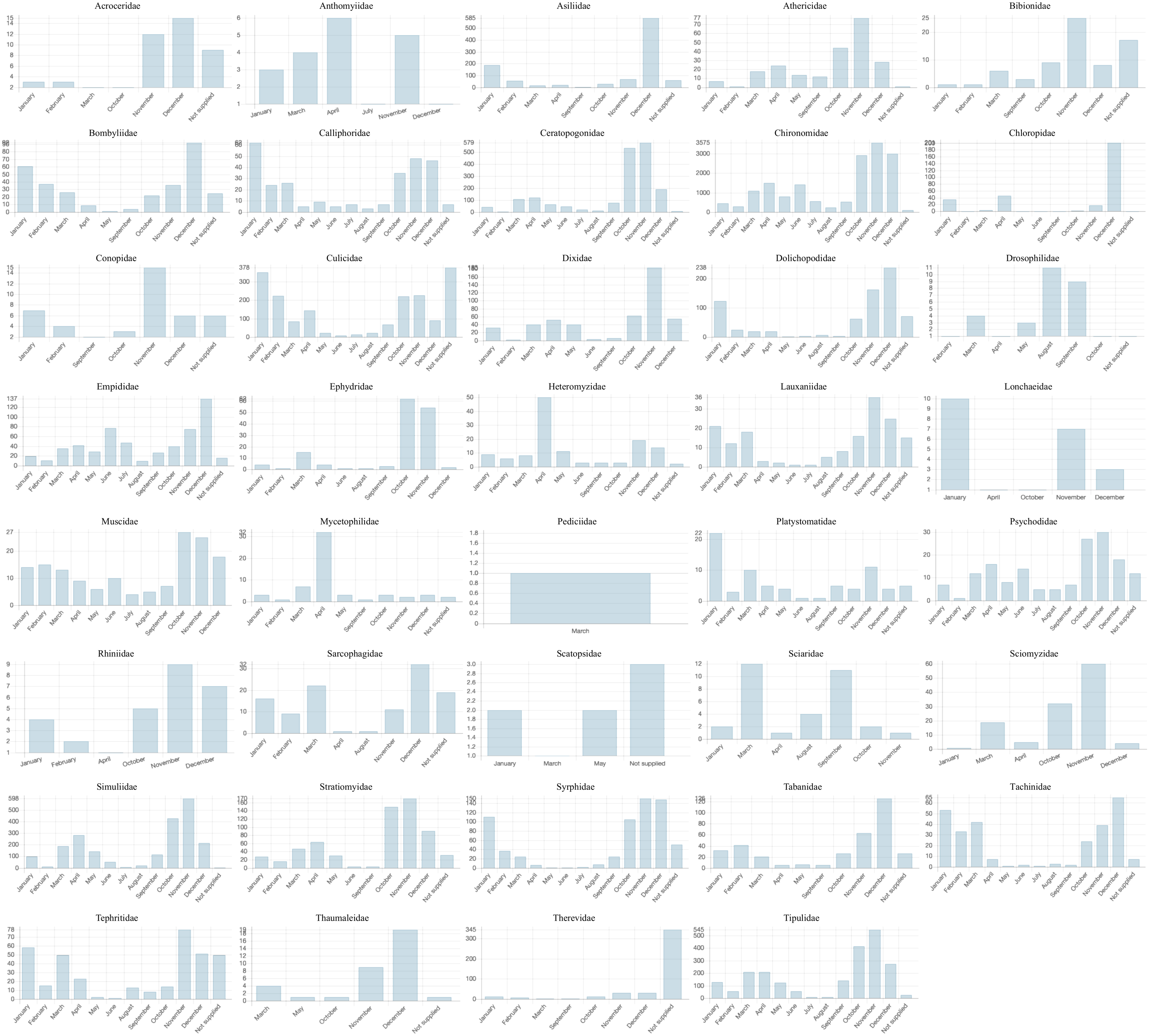

(b)

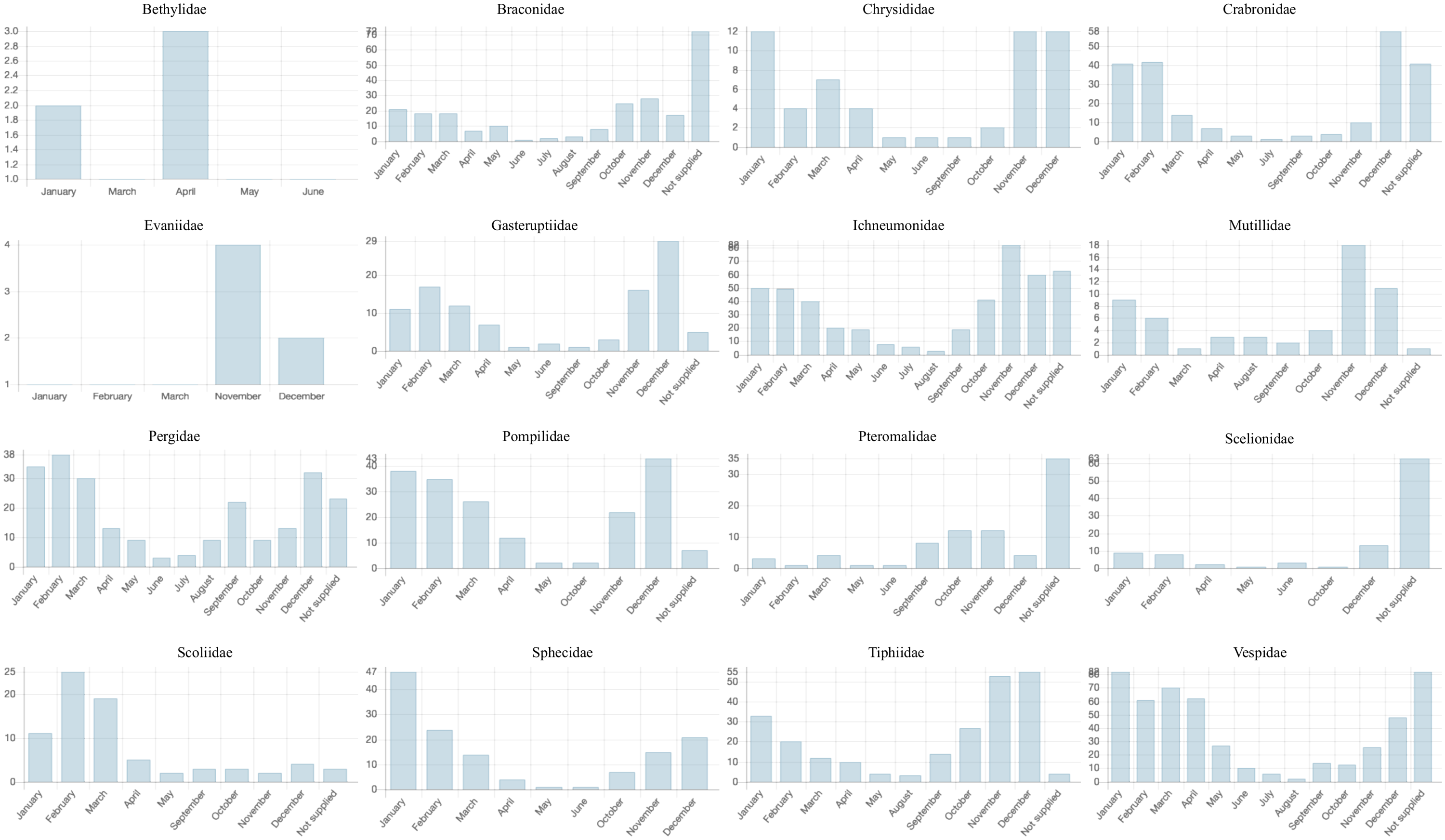
